# Dynamic metabolic profiles of marine macroalga *Ulva prolifera* during fragmentation-induced proliferation

**DOI:** 10.1101/578963

**Authors:** Yanli He, Yanhui Wang, Chaoyang Hu, Xue Sun, Yahe Li, Nianjun Xu

**Affiliations:** Key Laboratory of Applied Marine Biotechnology of Ministry of Education, School of Marine Science, Ningbo University, Ningbo 315211, China; Zhejiang Pharmaceutical College, Ningbo 315211, China

## Abstract

*Ulva prolifera*, a type of marine macroalgae, is the causative species behind green tides mainly in the Yellow Sea and adjacent regions, nevertheless, it can be used as food or animal feed in South China. The vegetative fragments of *U. prolifera* are an important seed source for successive green tide blooms. Fragmentation shortens the transition time from the vegetative state to the reproductive state. However, the translation of the algal metabolites during gametogenesis was far from well understood. In this study, the dynamic metabolic profiles of *U. prolifera* thallus during fragmentation-induced proliferation were investigated using non-targeted metabolomics approach in a time series of experiments in June 2017. After a 30 min low temperature shock, fragmentation induced a reproductive response of 91.57% of *U. prolifera* in 48 h, whereas that was only 21.43% in the control group. A total of 156 chromatographic peaks were detected, and 63 metabolites were significantly changed in *U. prolifera* during reproduction. The results of the Kinetic metabolic pattern showed that the fragments not only induced the formation of sporangium, but also complicated their metabolites accumulation. During fragmentation-induced proliferation, *U. prolifera* consumed different sugars at different time points. γ-aminobutyric acid (GABA), glutamic acid, gallic acid, and malic acid may play important roles in germ cell formation and release of *U. prolifera*, whereas n-hexanol, 2-methyl-3-phenylindole, and 3-indoleacetonitrile may be beneficial in biotic stress resistance. Compared with the control group, the metabolites, such as alcohol and organic acid, also showed significant difference with the photoperiod at the initial stage of proliferation (before 60 h). In conclusion, that the metabolites including sugars, organic acids, and alcohol changed with different photoperiod may be the strategy for *U. prolifera* to cope with adverse environment and rapid proliferation.

## Introduction

Marine macroalgae are potential sources of oil, food, and valuable compounds[1]. *Ulva* spp. are attractive models for understanding plant growth, development, and evolution, but they can cause severe ocean green tides. *Ulva* species generally have complex life cycles, which alternate between diploid and haploid generation, and exhibit parthenogenetic development of gametes. The germ cells of *Ulva* species are transformed directly from vegetative cells [2]. This transformation is caused by external factors. Tissue fragmentation is considered to be an important factor in inducing this transformation and reproduction quickly [3]. Fragments significantly increase the sporulation rate of *U. mutabilis* from 15.8% to 80.0% [4]. Within 2-3d after fragmentation, the reproduction of *U. pertusa* was induced [2]. Gao et al (2010). found that almost all *U. prolifera* fragments with a diameter of 0.5 mm were transformed into sporangium, whereas the sporangium in larger fragments was only formed by marginal and sub-marginal cells [5].

The mechanism of fragmentation-induction has been preliminarily considered that cutting destroyed cell walls and the additional cell-matrix structure, making it easier for algal segments to filter out inhibitors, thus removing the regulatory barriers to reproduction [2]. The gametogenesis of mature blades can be artificially induced by mincing the thallus into single mono-layered fragments and by washing-out of the sporulation inhibitors[6–8]. Previous studies showed that fragmentation-induction only improved the conversion rate of *Ulva* sporangium, and *Ulva* filaments can also be transformed into sporangium under the same culture conditions[4,5]. Cell differentiation is a popular topic in macroalga research[9,10]. However, the changes of the algal metabolites in space and time are far from well understood. Thus, we aimed to identify changes of the chemical composition of the metabolome during gametogenesis of *Ulva prolifera*. The metabolomics is a powerful tool for the study on the metabolism and physiological functions of marine organisms including seaweed [11,12]. Several metabolomic studies on the growth and reproduction of plants have been conducted[13]. In recent years, studies on marine macrophytes such as *Ectocarpus, Porphyra*, and *Zostera* by GC-MS(gas chromatography-mass spectrometry) have provided a comprehensive understanding of the metabolic networks linked to photorespiration pathways, tricarboxylic acid cycle, glycolysis, and pentose phosphate pathway involved in the acclimatization to external stressors[13,14]. In this study, the metabolites of *U. prolifera* in both natural reproduction and fragmentation-induced reproduction at different time points were investigated, and the results may help in exploring the causes of the rapid formation of marine green tides.

## Materials and methods

### Culture conditions

Vegetative *U. prolifera* thalli were collected from Xuwen Seaweed Co., Ltd., Xiangshan County, Zhejiang Province, China (121°56′153, 29°05′065), in June 2017. The thalli were gently washed three times with sterile seawater and then rinsed thoroughly. Using a magnifying glass, the attached sediments, small herbivores, and epiphytes were removed with a brush. Afterward, the thalli were cultured in f/2 medium with the following specific parameters: light intensity, 70 mmol photons m^−2^ s^−1^; temperature, 25 ± 1 °C; and light:dark cycle, 12 h:12 h in an intelligent illumination incubator (GXZ-280 C, Ningbo Jiangnan Instrument, China) at a salinity of 22 ± 1 for acclimation for about a month before the experiments. Ampicillin (100 mg.L^−1^), gentamycin (50 mg.L^−1^) and GeO_2_ (0.3 mg L^−1^) were added to suppress the growth of bacteria and diatoms[15] The medium was aerated and refreshed every other day. Unless otherwise specified, the following experiments were performed under the same conditions. The formal experiment began after 20 - 25 d of domestication.

### Sample treatments

Treatment group: *Ulva* thalli were harvested at morning and cut into small single-layered fragments (200 - 500 cells fragment) using a Zyliss ^®^ Smart Clean Food Chopper. Two grams of fresh algae fragments were cultured in a 3000 mL flask containing 2500 mL of sterile artificial seawater. Control groups: two grams of fresh uncut vegetative thalli was cultured under the same conditions. Both were pre-cultured at 4 °C for 30 min in a refrigerator and performed with six duplicates. Samples were collected at culture times of 12, 24, 36, 48, 60, 72, 96, and 120 h, and the media were reduced accordingly to maintain the same culture density.

### Microscopic observations and counting methods

The formation and release of *U. prolifera* germ cells were observed under an AOTE BK5000 microscope equipped with a Cool SNAP digital camera system. Seven microscopic fields of view were randomly observed in each selected fragments or uncut thalli. During the maturation process, the germ cell sporangium and released germ cell sporangium were imaged to count their numbers. The percentage of mature germ cell sporangium and the formation and release rates were calculated using the following equations: Sp = (Sm /At)×100%, Rp = (Ss/At)×100%,where S_p_ represents the percentage of mature sporangium, S_m_ represents the mature sporangium, R_p_ represents the percentage of released germ cell cysts per unit of thalli, S_s_ represents the germ cell cysts that were released (in μm^2^), and At represents the number of germ cell cysts in the whole field under the microscope.

### Metabolite extraction and GC-MS analysis

#### Extraction

Sample materials (20mg±2mg) were put in 2mL Eppendorf tube. One metal or zirconia ball was added in every tube. 700μL 100% methanol and 5.75 μL Stock ribitol (Sigma-Aldrich, München, Germany, 0.2 mg/mL Stock in Water) were added, then vortexed for 15 seconds. Following by homogenized with the Retschmühle for 3 min at 25 Hz. Centrifuged for 10 min at 14000 rpm and take 600 μL from the upper phase into 2mL fresh Eppendorf tubes, then 300μL CHCl_3_ added and mixed it slowly. After that, 750 μL H_2_O was added in and vortex 15 sec. Subsequently, centrifuged 10 min at 14000 rpm and 150 μL upper phase (polar phase) was taken into 1.5mL Eppendorf tubes. Finally, all the extracts were dried in a speed vac for at least 3 h in normal temperature condition.

#### Derivatization

After evaporation of the solvent, 40 μL of methoxyamine hydrochloride (Sigma-Aldrich) solution (20 mg/mL) in pyridine was added for derivatization and incubated at 60°C for 1 h. Subsequently, 50 μL of N-Methyl-N-(trimethylsilyl) trifluoroacetamide (Macherey-Nagel, Düren, Germany) was added to each sample and incubated at 40 °C for 1 h, then transferred 100 μL into sample vials and analyzed directly by GC-MS.

#### GC-MS analysis

The metabolites were detected using the Trace 1300/ITQ 900 GC-MS system (Thermo Fisher Scientific, USA, Serial No.13005). GC-MS analysis were conducted as previously reported [16] with slight modifications. The samples were injected with a 1300 autosampler equipped with a 10-μL tapered, fixed needle, polytetrafluorethylene (PTFE)-tipped plunger syringe. The column and all spare parts were purchased from Thermo Fisher Scientific (Thermo Fisher Scientific, USA). The GC parameters for the analysis were as follows: The inlet temperature was 250°C, EI ion source temperature was 230°C, high-purity helium (99.99%) as the carrier gas, an amount of 1 μL sample was injected with the splitless mode. The temperature was initially set at 60 °C for 2 min, then increased by 10 °C min^−1^ up to 150 °C and held for10 min. Then, the temperature rised to 250 °C at 15 min^−1^ and held for 20 min. MS was performed with a full-scan mode with MS range from 0 to 550 (m/z).

#### Data analysis

Compounds were identified based on the ratio and relative abundance of the National Institute of Standards and Technology database (NIST) and MassBank website (http://www.massbank.jp/) according to mass/charge. Statistical analyses were performed in MetaboAnalyst 3.0 (http://www.metaboanalyst.ca/) according to its protocol (Linting and Meulman 2011). “Normalization by internal standard ribitol” was chosen for sample normalization, “None” for data transformation, and “Auto scaling” for data scaling. Principal component analysis (PCA) and orthogonal partial least squares discriminant analysis (OPLS-DA) were performed in the SIMCA-P 13.0 software package using Auto scaling. Two-sided T-test, one-way ANOVA, two-way ANOVA, post-hoc analysis, and ANOVA simultaneous component analysis (ASCA) were performed in MetaboAnalyst 3.0 (http://www.metaboanalyst.ca/) according to its protocol[17]. “Auto scaling” was selected as the data scaling method for all analyses. The two-way ANOVA type used was “within subject ANOVA.” Statistical significance was considered at *P* < 0.05, and the false discovery rate was used for multiple testing correction. For ASCA, the alpha and leverage thresholds were set as 0.05 and 0.9, respectively. Figures were edited with Adobe Illustrator CS5 for better resolution.

## Results

### Microscopic observation of the germ cell sacs formation

As shown in Fig. 1, the mature germ cell sac first appeared on the edge of the cut algae at 24 h, but most of the cells remained in a vegetative state at 60 h in the control group. At 48 h, nearly all of the vegetative cells (91.57%) (Table 1) in the treatment group were transformed into germ cell sacs, and the germ cell sacs that matured early at the edge began to release. However, only 9.57% of the vegetative cell transformation germ cell sacs until 60 h in the control group (Table 1). At 72 h, nearly all of the vegetative cells (98.29%) were transformed into mature germ cell sacs in the treatment group, meanwhile, 20.43% of the algae were transformed into germ cells (Table 1) in the control group. At 96 h, almost all of the cells were transformed into mature germ cell sacs. Moreover, germinated seedlings appeared in the treatment group. However, only 4.28% germ cell sacs were released in the control group although most of the vegetative cells were transformed into germ cell sacs (120 h, Fig. 1).

**Fig. 1.**
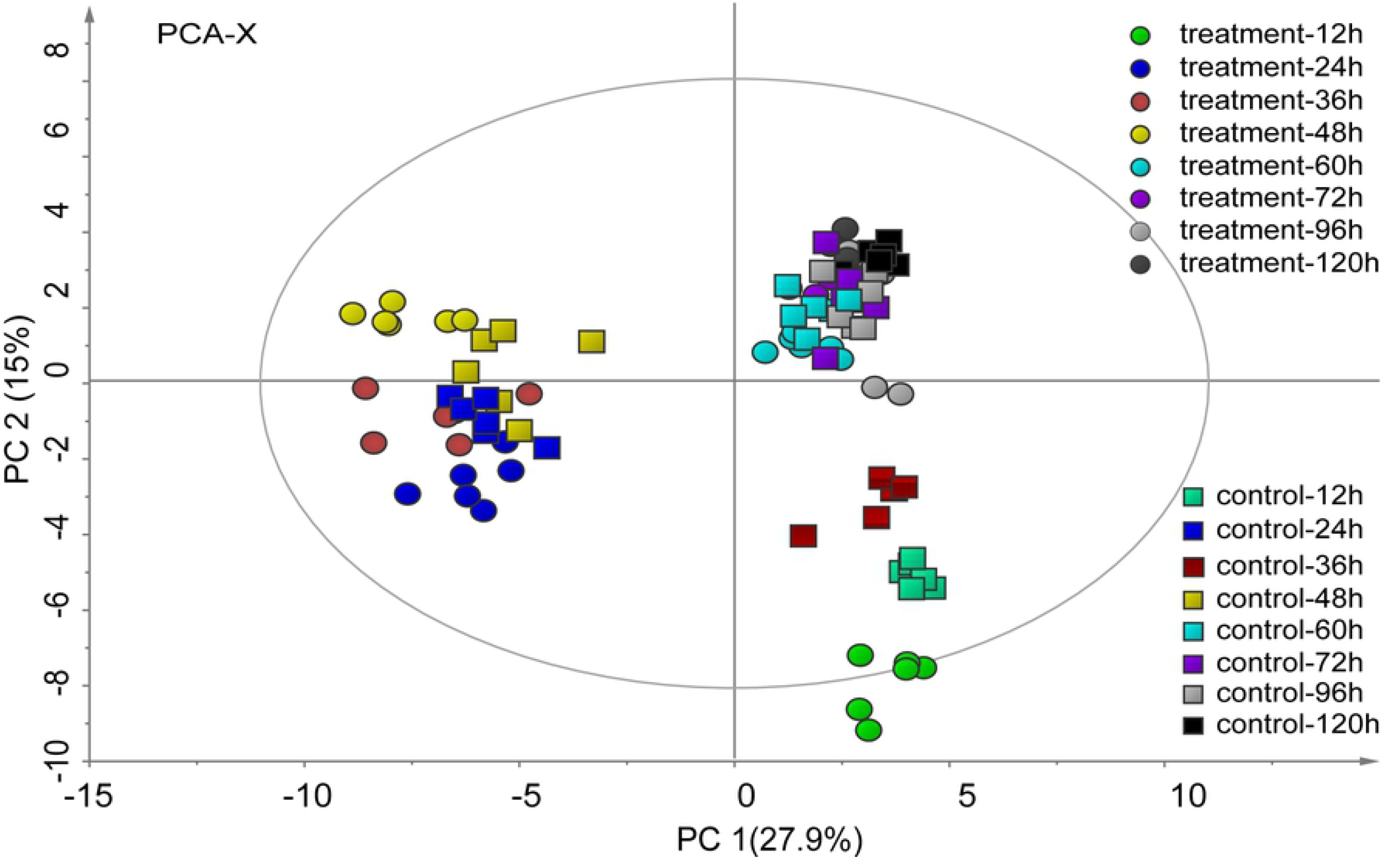
Microscopic observation of the transformation of *U. prolifera* from the vegetative state to the reproductive state after cutting (left: uncut(control); right: cut (treatment) 12h, 24h, 36h, 48h, 60h, 72h, 96h, 120h) (in treatment 48h: a, b, c, d, represent vegetative cell; mature germ cell sac; released germ cell sac; and released gametophytes/sporophytes respectively; in treatment 120h: e represents germinated seedling), the scale bars represent 20 μm.

**Table 1**. Transformation of *U. prolifera* from vegetative cells into germ cell sac and their release rate.

^a^ percentage of mature germ cell sporangium

^b^ release rates of germ cell sporangium

^c^ average of six tests

### Analysis of metabolite profiles

To investigate the metabolic profiles of *U. prolifera* during fragmentation-induced proliferation, both the filament and fragments at 12, 24, 36, 48, 60, 72, 96, and 120 h were collected and subjected to metabolic profiling analysis based on GC-MS. Smoothing and baseline correction was performed prior statistical analysis. After the background subtraction, the impure peaks due to column material loss or impurity caused by the sample preparation process were removed, only m/z values with signal-to-noise ratio more than 10 were picked out. As a result, 156 chromatographic peaks were detected. The GC-MS data were eventually organized into two-dimensional matrices, including observed values (samples) and variables (peak intensities). PCA was subsequently performed on the 156 metabolites to obtain overall patterns within the kinetic metabolic patterns and induction methods of reproducing *U. prolifera*. The first principal component (PC1), accounting for 27.9% of the total variance, reflected time-dependent *U. prolifera* reproduction, and samples from treatment and control groups were gathered at the same time point (Fig. 2). The results indicated that the developmental time affected *U. prolifera* metabolites more than the induction method did. The second principal component (PC2) accounted for 15% of the total variance. The samples of the treatment and control groups at the same time points were all distinguishable in PCA plots, indicating that the metabolites with different treatments of *U. prolifera* were different at the same developmental times.

**Fig. 2.**
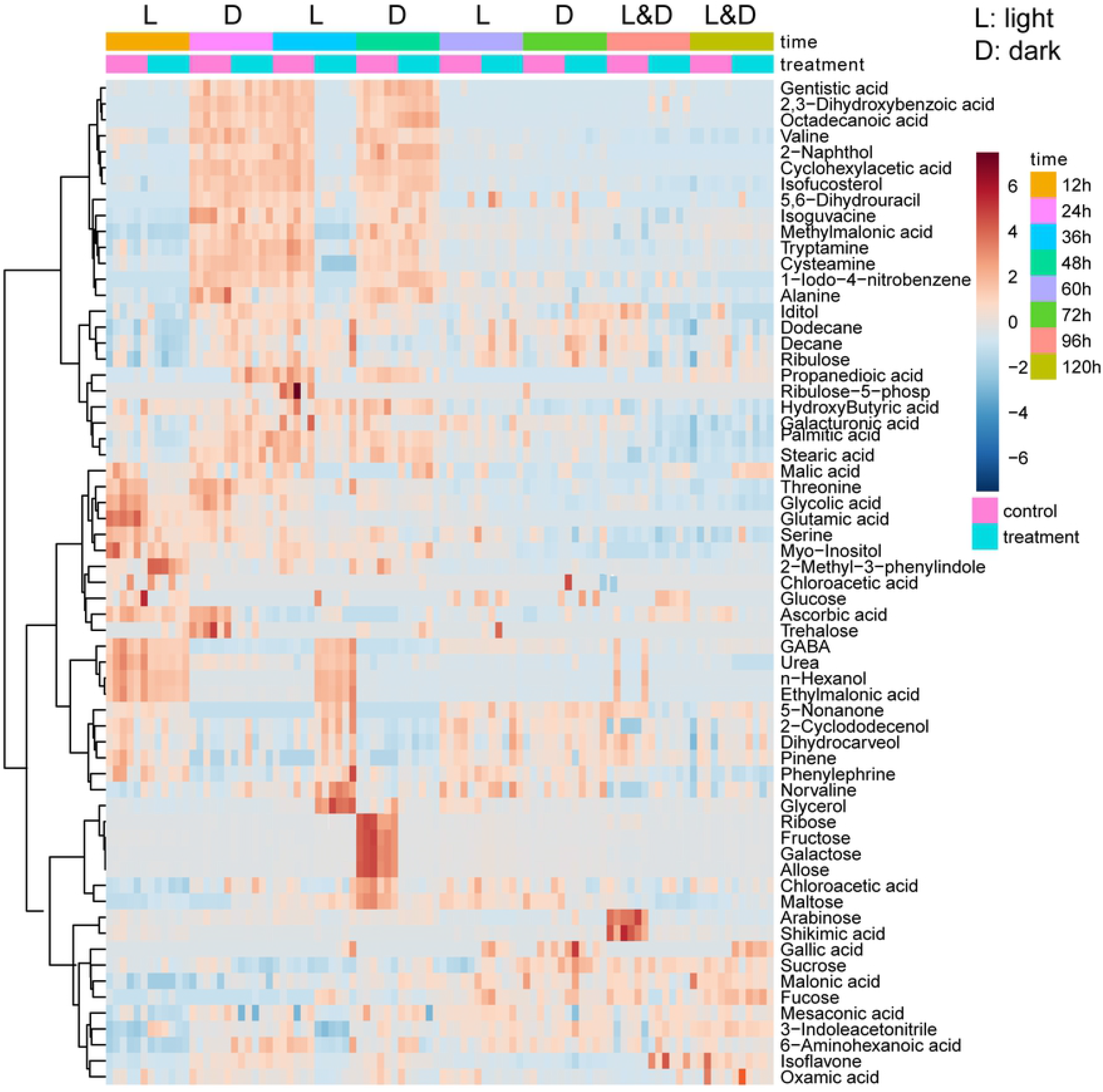
PCA of *U. prolifera* during its fragmentation-induced proliferation (Colors represent different treatment times, boxes (□) represents the control group, and cycle (○) represents the treatment group; n=6).

PCA provides a high-level summary of the main patterns of data variance. Detailed metabolite abundance profiles can be obtained from the visualization of metabolomic data based on heat map. The changes of metabolites in the cut and uncut thallus over time are mainly the same. Of note, these metabolites exhibit opposite trends at the 36 and 48 h time points with respect to the treatment and control groups (Fig. 3).

**Fig. 3.**
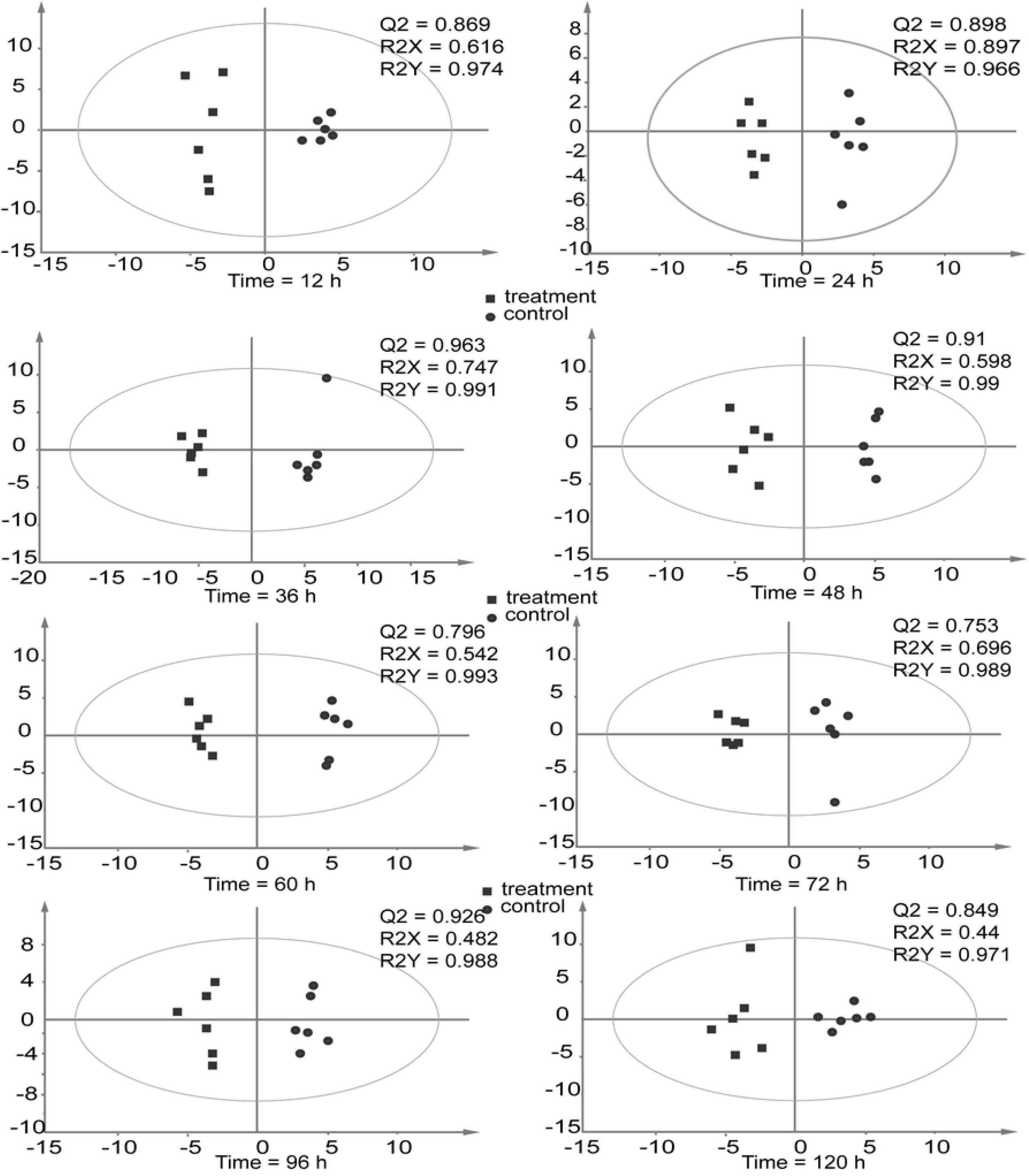
Heatmap analysis, combined with HCA, of the metabolites in *U. prolifera* fragmentation-induced proliferation (treatment and control groups) during 12-120 h (n=6).

Two-way repeated measures (within subjects) ANOVA was then utilized to analyze which factors (treatment, developmental time, and their interaction) caused the variation of each metabolite. The abundance of 63, 32, and 58 metabolites was affected by treatment, developmental time, and their interaction, respectively (Fig. 4).

**Fig. 4.**
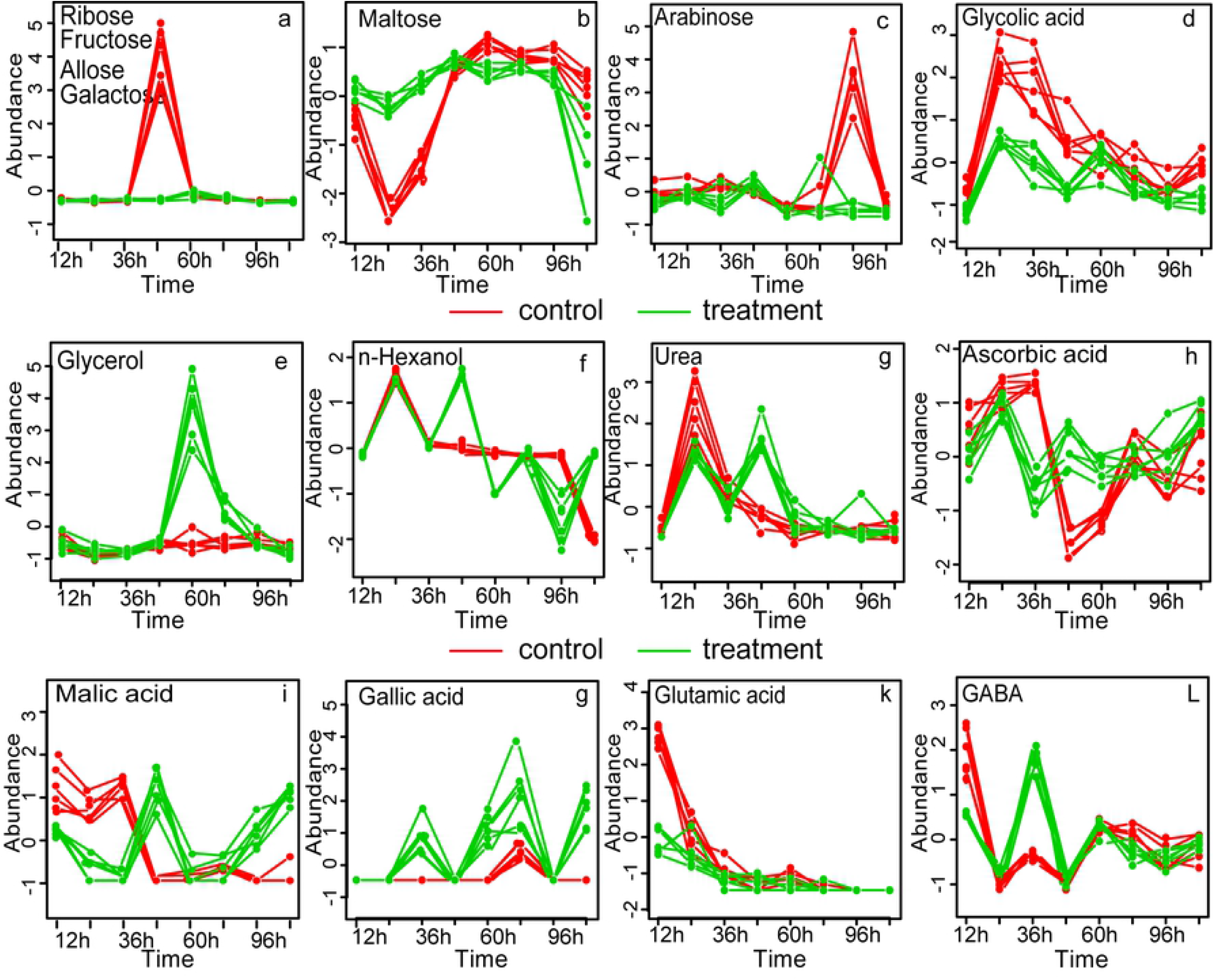
Two-way repeated measures (within subjects) ANOVA of metabolites in *U. prolifera* fragmentation-induced proliferation during 12-120 h.

ASCA was performed to determine the trends of treatment methods, developmental time, and their interaction patterns[17]. The time score plots based on component 1 (45.21% of variation explained) of the corresponding model demonstrated a decrease in scores from 12 h to 36 h, then increased at 48 h, then decreased from 48 h to 60 h, and final increased until 120 h (Fig. 5a). The treatment score plot showed that different dispose types differed in their PC1 scores, with the treatment and control groups having the lowest and highest scores, respectively (Fig. 5b). Component 1 of the interaction effect obviously exhibited opposite trends at 48 h between the treatment and control groups, while the same trend was observed at other times (Fig. 5c).

**Fig. 5.**
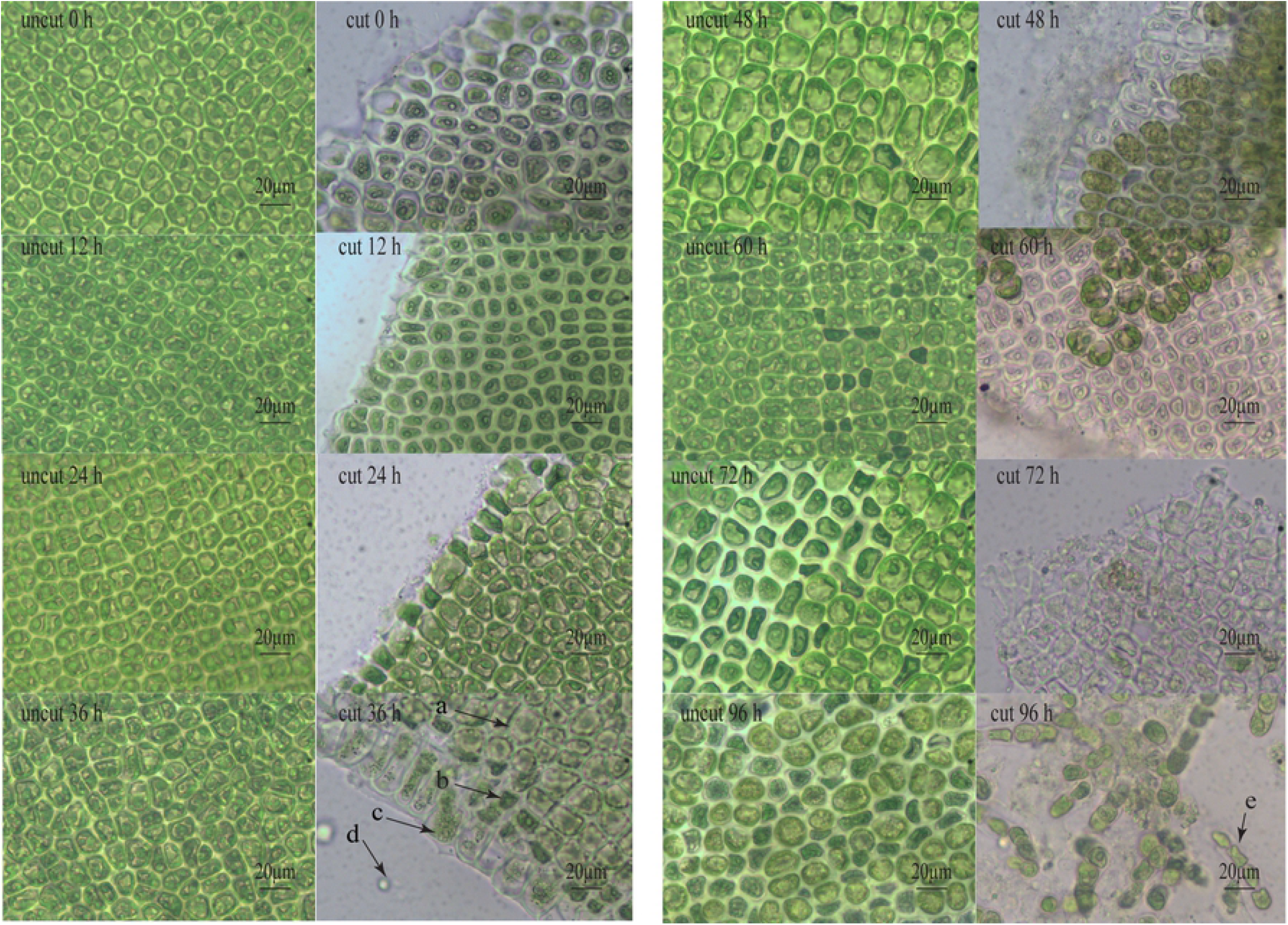
ANOVA-simultaneous component analysis (ASCA) of *U. prolifera* during its fragmentation-induced proliferation.

Leverage/squared prediction error (SPE) plots were made to correlate the metabolic features with the experimental factors[17]. The leverage evaluates the importance of the metabolite to the model, and SPE tests the fitness of the model for particular metabolites. Metabolites with high leverage SPE, which contribute significantly to the model, were chosen as well-modeled metabolites. In this way, 63 metabolites which were affected by treatment were analyze by SPE, a total of 31 metabolites were well modelled, including 3 alcohols, 5 amino acids, 11 organic acids 6 sugars and 6 other kinds metabolites (Table S1). Twelve well-modeled metabolites, including six organic acids (cyclohexylacetic acid, gentisic acid, octadecanoic acid, 2,3-dihydroxybenzoic acid, stearic acid, palmitic acid), two kinds of alcohol (isofucosterol and 2-naphthol), one amino acid (valine) and two other kinds of metabolites (tryptamine and 5-nonanone), were determined based on the major pattern of time (Fig. 5d and Supplementary Table S1). Their levels sharply declined from 0 h to 24 h in the filament and its fragments. Then, they increased at 36 h, decreased at 48 h, and peaked at 96 h. Eight metabolites, including glycolic acid, glycerol, alanine, valine, arabinose, norvaline, chloroacetic acid allyl ester, and ribose, were well modeled by treatment method (Fig. 5e and Supplementary Table S1). Eleven metabolites, including fructose, allose, ribose, glycerol, cysteamine, galactose, ribulose-5-phosphate, ascorbic acid, 3-indoleacetonitrile, methylmalonic acid, and malic acid, were well modelled (Fig. 5f and Supplementary Table S1) based on the interaction, and their levels varied between the filament and its fragments.

### Metabolic difference analysis

To simplify the information and obtain maximal covariance among metabolite levels during proliferation at different time points, OPLS-DA was applied. The OPLS-DA model, which was composed of one orthogonal component and one prediction component, generated the explained variation values R2X (cum) > 0.82 (except at 96 and 120 h where the R2X values were 0.482 and 0.44) and R2Y (cum) > 0.966 and the predictive capability Q2 (cum) > 0.753 at all six time points (Fig. 6). These high-value parameters indicated the excellence in modeling and prediction with good discrimination between the fragmentation-induced proliferation and natural proliferation of *U. prolifera* at each time point before 96 h because the model parameters (>0.5) are considered to be satisfactory in explanatory and predictive capabilities [18].

**Fig. 6.**
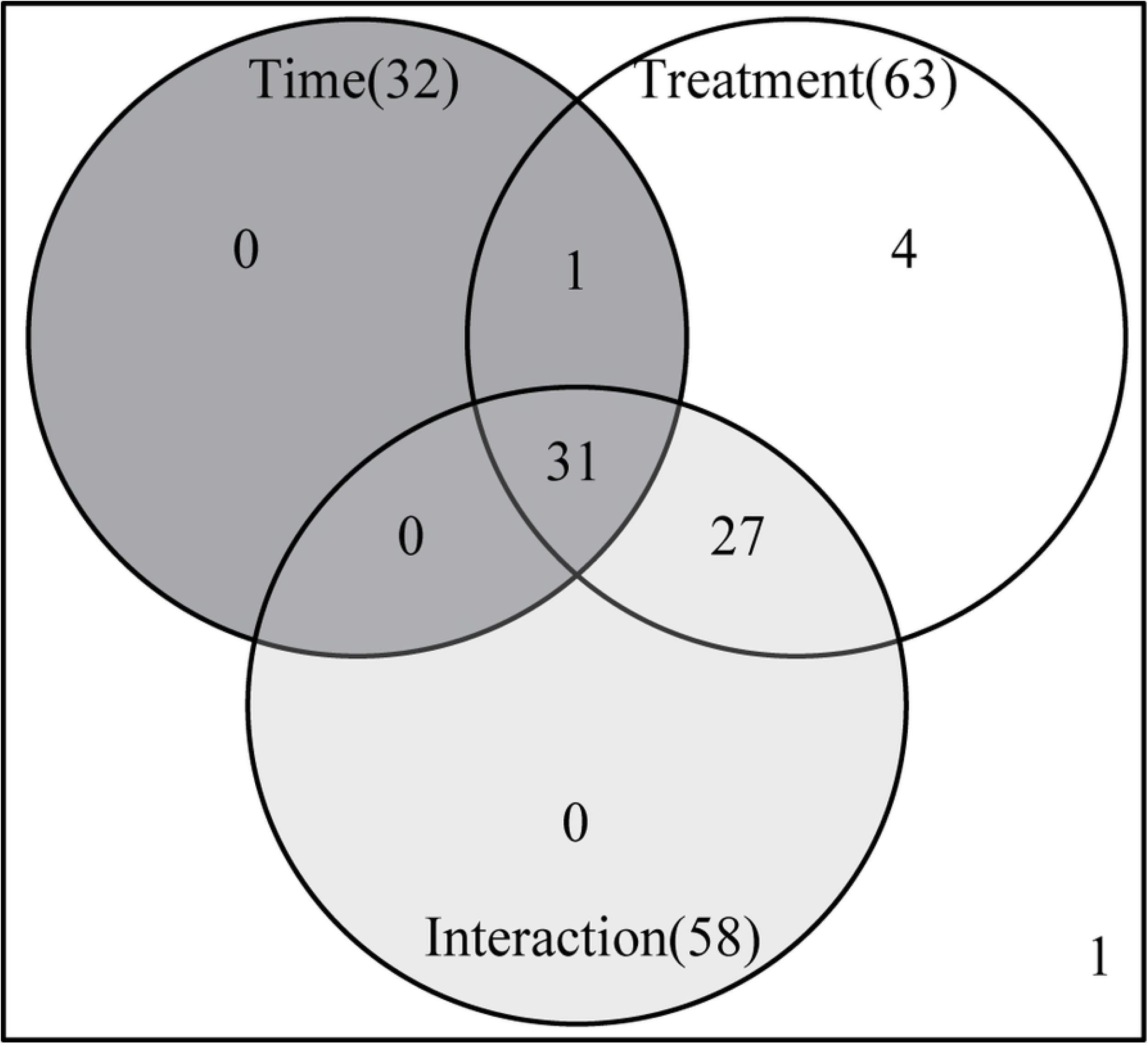
OPLS-DA of the uncut and cut treatment of *U. prolifera* at different time points.

Based on a variable importance in projection (VIP) threshold (VIP > 1) from the 200-fold cross-validated OPLS-DA models, a number of metabolites associated with the distinction of metabolic profiles of fragmentation-induced proliferation and natural proliferation and fragmentation-induced proliferation at each time point were procured. The candidate metabolites from the OPLS-DA model, whose P-values are less than 0.05 between fragmentation-induced proliferation and natural proliferation at each time point tested using the T-test with fold threshold of > 2 or < 0.5, were further selected as biomarkers (Table 2).

**Table 2**. Metabolic difference between cut and uncut groups of *U. prolifera* at each time point.

^a^ P-value was calculated from T-test (two-tails) with a cutoff of 0.05.

^b^ Ratio of relative metabolites at 24 and 12 h after cut.

^c^ VIP was obtained from OPLS-DA with a threshold of 1.0.

^d^ Comparison of relative content of metabolites in *U. prolifera* at each time point.

At 12 h, four metabolites, maltose, 2-methyl-3-phenylindole, 3-indoleacetonitrile, and glutamic acid, were selected as biomarkers (Table 2). The levels of maltose (12.23-fold), 2-methyl-3-phenylindole (2.78-fold), and 3-indoleacetonitrile (2.33-fold) were significantly higher and the relative abundance of glutamic acid was significantly lower than that in the control group.

At 24 h, three metabolites, maltose, propanedioic acid ester, and trehalose, were important in distinguishing between the treatment and control groups (Table 2). In the treatment group, the relative contents of trehalose significantly decreased, however, the relative content of maltose increased by 12.44-fold, and those of propanedioic acid ester also significantly increased.

At 36 h, a total of 18 metabolites, including six kinds of acids, two kinds of alcohol, one benzene derivative, one kind of lipid, six flavonoids, three lipids, two carbohydrates, and six other kinds of metabolites, played important roles in separating the treated and control groups. In the treatment group, the top five metabolites with the highest fold values were n-hexanol, 5-nonanone, GABA, ethylmalonic acid, and pinene. In particular, the relative contents of n-hexanol was 15.09-fold higher than that of the control group. The levels of five metabolites, 1-iodo-4-nitrobenzene, gentisic acid, succinic acid, 2-naphthol, and cysteamine, were significantly decreased in the control groups, with cysteamine having the lowest level (0.09-fold).

At 48 h, eight metabolites, including six carbohydrates and two acids, were selected as biomarkers (Table 2). Among them, the relative content of four kinds of monosaccharides (galactose, ribose, allose, and fructose) and glycerol decreased by 0.39–0.03-fold compared with those of the control groups. The levels of malic acid and ascorbic acid increased significantly by 9.61- and 6.05-fold, respectively. However, the levels of maltose, glycerol, allose, galactose, ribose, and fructose decreased.

At 60 h, three metabolites, including glycerol, arabinose, and glutamic acid, changed significantly between the treated and control group. Their relative contents were all decreased in the fragmentation-induced proliferation groups. At 72 h, only two metabolites (maltose and n-hexanol) were selected as biomarkers.

However, at 90 h, four metabolites, malic acid, norvaline, isoflavone, and glucose, accumulated significantly in the treatment group. Meanwhile, the relative contents of seven kinds of metabolites, including arabinose (0.11-fold), maltose (0.14-fold), ribose (0.26-fold), fucose (0.42-fold), shikimic acid (0.09-fold), iditol (0.27-fold), and 5-nonanone (0.29-fold), sharply decreased.

At 120 h, malic acid and gallic acid increased, and urea decreased compared with those in the control group.

### Kinetic metabolic pattern

In order to investigate the variation of the 31 metabolites well modelled by SPE, multivariate Empirical Bayes Analysis (MEBA), a time-course analysis method based on the MEBA statistic, which can evaluate the importance of features by Hotelling’s T, was employed to identify the metabolites, with differential temporal profiles [19]. Twenty-three metabolites, which meet the following conditions: potential biomarker (Table 2), the Hotelling’s T2 value > 100 and well-modelled according the analysis by ASCA are shown in Fig. 7 (n-hexanol, cyclohexylacetic acid, alanine and urea varied obviously with time but did not meet the above three conditions at the same time) (Table 3). The changes of four carbohydrate metabolites (ribose, fructose, allose, and galactose) with time were consistent: from 0 h to 36 h, their abundance varied between −1 and 0, and almost reached 4 at 48 h in the control group. Then, the abundance sharply decreased to nearly 0 at 72 h, while that of the treatment group was relatively stable (Fig. 7a and Table 3). This result was consistent with the heat map. Take maltose for example, its abundance in the control group decreased from 0 to −2.5 and then gradually increased until reaching the highest at 60 h, followed by slight fluctuation with the photoperiod and maintaining a value between 0 and 1 at 0–48 h. The change trend was the same as that in the treatment group. However, the abundance of maltose in the treatment group fluctuated slightly between 0 and 1 with the photoperiod at 48–96 h and then decreased to below 0 at 96–120 h (Fig. 7b and Table 3). In the treatment group, the relative content of the four alcohols was related to the photoperiod before 60h, and the relative content increased at night (without light) and decreased during the day (with light), after 60 h, the relative content gradually leveled off. In the control group, however, 36 h was a cutoff point, where the relative content increased before 36 h and then decreased gradually (except for glycolic acid, where the relative content decreased gradually from 12 h). Interestingly, the relative content of other six metabolites including three organic acids (gentistic acid, ascorbic acid and cyclohexyacetic acid), valine, 1-Iodo-4 nitrobenzen and cysteamine also showed this trend.

**Fig. 7.**
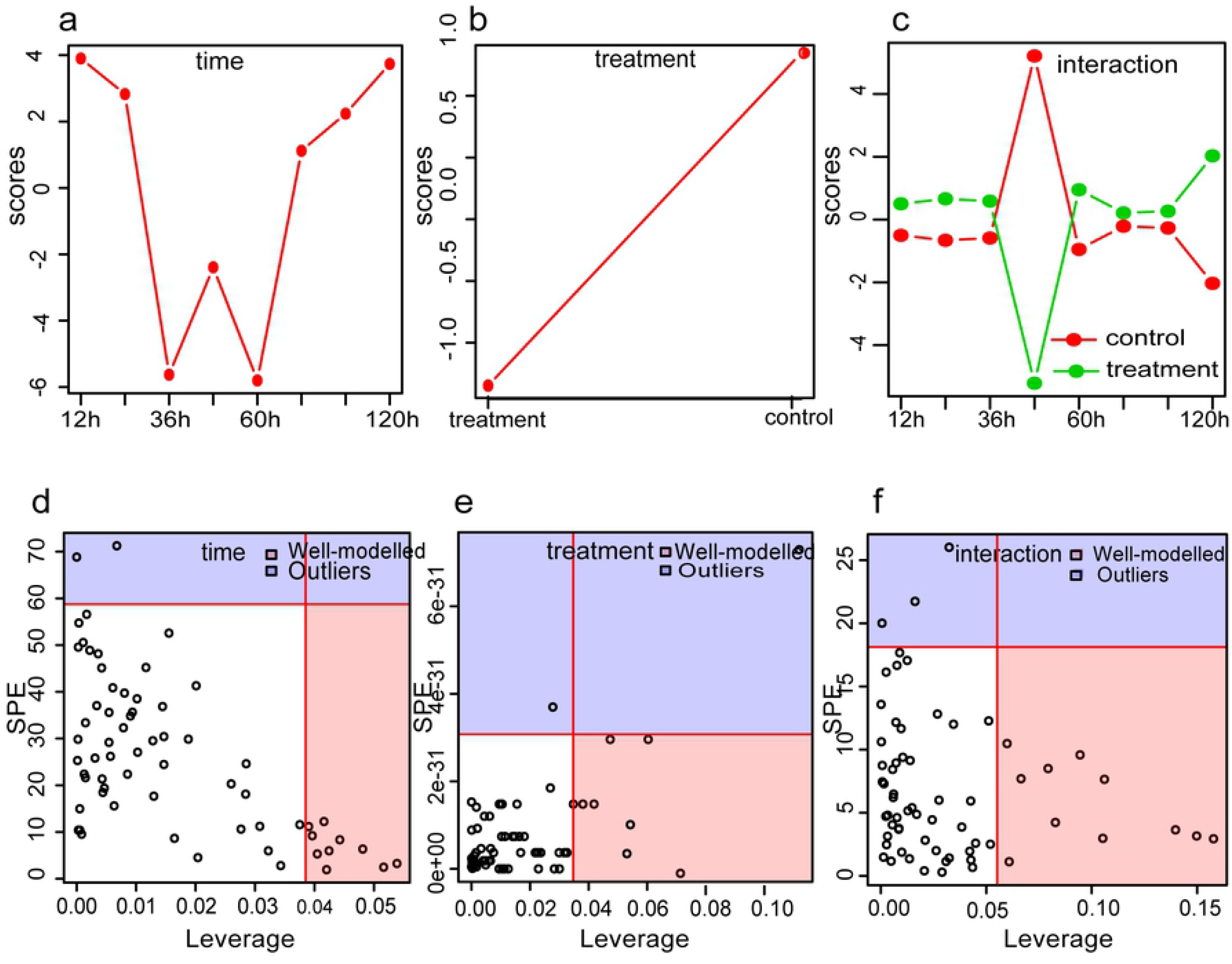
Metabolites MEBA of *U. prolifera* which meet following conditions: potential biomarkers according OPLS-DA, well-modelled by SPE and the Hotelling’s T2 value > 100.

**Table 3** Metabolites of *U. prolifera* which meet following conditions: potential biomarkers according OPLS-DA, well-modelled by SPE and the Hotelling’s T2 value > 100.

## Discussion

Fragmentation-induced spores play a vital role in the rapid accumulation of large amounts of biomass and may be one of the most important factors in accelerating the occurrence of green tides on the coast of Qingdao China [5]. In this study, reproductive cells (nearly 90%) (Table1) appeared within 48 h after *U. prolifera* thalli were cut into approximately 0.5 mm segments. It was consistent with the results of the study on *U. mutabilis* [4]. Interestingly, almost all the cells (96.29%) in the physiological state transformed into the reproductive state at 120 h in the control group. It indicated that fragmentation-induction was not the determinant of *U. prolifera* sporangium formation, but only a promoter. The metabolic profiles of the treatment group differed significantly from that of the control group according to PCA and OPLS-DA. Time-course analysis showed that the relative content trend of carbohydrates, alcohols and organic acids was different in treatment group and control group (Fig 7).

### Different sugars response during germ cell formation

Sugar is mainly involved in the biosynthesis of sugar polymers, including starch and cellulose, generation of energy (ATP), cross-talk in invertase-mediated sugar signaling, and phytohormone control [20]. The content of monosaccharides in the treatment group was significantly lower than that in the control group at two time points: 48 h (ribose, fructose, allose, and galactose) and 96 h (arabinose) (Fig. 3 and 6), the results were consistent with the kinetic metabolic pattern analysis (Fig.7a). Meanwhile, the content of maltose was higher in the treatment group within 12 - 48 h after cutting and then became similar to that in the control group (Fig.7b). The result indicated that in the process of fragmentation-induction and natural proliferation, *U. prolifera* consumed different sugars. Kaplan and Guy (2004) indicated that the resultant maltose accumulation may function as a compatible-solute stabilizing factor in the chloroplast stroma in response to acute temperature stress [21]. In this study, the accumulation of maltose suggests that it was not only related to energy metabolism during the spore formation of *U. prolifera*, but also related to the response to mechanical injury.

### Organic acids may be involved in the redox function and osmotic regulation of *U. prolifera*

Compared with the control group, several organic acids accumulated, including ethylmalonic acid (4.35-fold) at 24 h, malic acid (9.61-fold) and ascorbic acid (6.06-fold) at 48 h, and gallic acid (7.07-fold) and malic acid (7.02-fold) at 120 h (Table 2). During the process of transformation from vegetative to reproductive state, the pigmentation deepens, the vacuole becomes smaller and then disappeared at treatment 36 h and control 48-60 h (Fig.1). The accumulation of organic acids may contribute to the regulation of the cell osmotic pressure to meet the need of the change from vegetative to reproductive state. Likewise, Alsufyani et al (2017) investigated that the composition of the metabolites released during Ulva grew under normal conditions with an exo-metabolomic approach and found that higher concentrations of divalent low-molecular metabolites such as maleic and succinic acids were detected in the late gametogenesis phase. An elevated level of organic acid is important for energy production, cellular membrane stabilization, maintenance of turgor, vitrification of cytoplasm and signaling in cells under stress conditions[13,22]. Moreover, ascorbic acid is one of the major antioxidant compounds synthesized in plants and used for cell division regulation [23,24]. In orthodox seeds, there is no ascorbic acid and ascorbic acid peroxidase activity in the stationary phase, but they restart after from the onset of imbibition[25]. Many studies have shown that ascorbic acid biosynthesis and rapid restart of ascorbic acid peroxidase activity are the keys to seed germination[26,27]. However, other studies have shown that high doses of ascorbic acid can inhibit the germination of wheat seeds [28]. In this study, 21.43% germ cells were formed at 36h, and 91.57% germ cells were formed at 48 h, and the germ cells started to release in the treatment group. In the meantime, the relative content of ascorbic acid decreased (0.24-fold, at 36h point) and then increased (6.06-fold, at 48h point) compared with those of the control group. In contrast to higher plants, reduced ascorbic acid content in *U. prolifera* may contribute to the formation of germ cells, and higher concentrations of ascorbic acid may contribute to the release of germ cells.

Accumulation of GABA is believed to be a redox regulatory mechanism in a variety of terrestrial plants, such as *Arabidopsis* [29], *Lotus japonicus* [30], *Pisum sativum* [29,31] and *Glycine max*[32]. Gallic acid and its related compounds showed significant chromosomal influence characteristics [33,34]. In this study, the relative content of GABA and gallic acid accumulated at 36h in the treatment group, the result was consistent with those of Singh et al.(2016), indicating that these two organic acids played a consistent role in *U. prolifera* and higher plants[23,33–36]

### The other metabolites in the formation of germ cells in *U. prolifera*

Compounds constitute phytochemical defense weapons from a variety of metabolic pathways and can be broadly divided into three categories, namely, isoprenoids (e.g., diterpenes), alkaloids (e.g., indole alkaloid camalexin), and shikimates (e.g., flavonoids) [37]. In this study, at 12 h, 2-methyl-3-phenylindole and 3-indoleacetonitrile accumulated in the treatment group (Table 2), suggesting that these metabolites were used for biological defense of *U. prolifera*.

Two obvious changes were noted in alcohol metabolites: glycerol and hexanol (Table 2). The content of glycerol in the treatment group was significantly higher than that in the control group at 60 h after cutting (Fig. 7e). In this study, the Sp of *U. prolifera* was 97% and the Rp was 9.57% at 60 h after cutting, but the they were only 32.86% and zero at 60h in the control group, respectively. Alsufyani et al. (2017) found that the maximum content of glycerol on the surface of *Ulva* was at 21h after artificial induction and the glycerol secreted by *U. prolifera* might be the carbon source of its epiphytes. In this experiment, the relative content of glycerol increased significantly at 60h (Fig.7E), which might be the result of the interaction between *U. prolifera* and the epiphytes which was beneficial for the formation of the germ cells[38]. Hexanol, a green leaf volatile that can be used as a synergist for insect pheromones, is used for controlling insect populations. In this study, the content of hexanol was not only affected by cutting, but also by photoperiod. The content of hexanol at night (24 h, 48 h, 72 h) was significantly higher than that in the daytime (12 h, 36 h, 60 h), indicating that hexanol might protect *U. prolifera* from nocturnal insect’s harm (Fig. 7f). Certainly, further targeted screening for organic acid is necessary to understand the functions of fragmentation-induced proliferation.

In conclusion, we found that, after cutting, the reproductive sporangia formed earlier than that in the control group and various of metabolites undergo drastic changes. The kinds of consumed sugars differ significantly at different time points during the formation of sporangium. At the initial stage of proliferation (the first 60 h), the metabolites such as alcohol and organic acid showed significant changes with the photoperiod, which may be the main strategy for *U. prolifera* to cope with adverse environment and rapid proliferation. However, GC-MS approach provides a very preliminary view of the metabolic changes that are the consequences or the causes of the differentiation of the reproductive cells. Identifying effective biomarkers to monitor complex cellular processes in space and time is only the first step before the challenging isolation and structure elucidation of the compound have to be performed in further studies. Besides, bacterial morphogenetic compounds might not be among them, as the biologically active concentration [39–41] seems to be far below the detection limit of the metabolomic approach applied [38]. As (epiphytic) bacteria are essential for *Ulva’s* morphogenesis, bacterial compounds are certainly part of the surface-associated metabolome. Future studies will explore these compounds by comparison, e.g., xenic *Ulva* cultures with normally developed adult axenic cultures grown in the presence of purified morphogenetic bacterial compounds[38].

## Acknowledgements

We thank the efforts of all people involved in the laboratory work and data analysis, and the collaboration of Dr Wenrong Zhu and Mr. Wei Hu for the donation of *Ulva* samples.

## Authors’ Contributions

N.X., C.H and Y.H. designed and supervised the research. Y.H. performed experiments, analyzed the data, and wrote the manuscript. Y.W. performed the GC-MS data analysis. All the authors revised and approved the final manuscript.

